# Convolutional neural network for automated mass segmentation in mammography

**DOI:** 10.1101/2020.12.01.406975

**Authors:** Dina Abdelhafiz, Jinbo Bi, Reda Ammar, Clifford Yang, Sheida Nabavi

## Abstract

**Background:** Automatic segmentation and localization of lesions in mammogram (MG) images are challenging even with employing advanced methods such as deep learning (DL) methods. We developed a new model based on the architecture of the semantic segmentation U-Net model to precisely segment mass lesions in MG images. The proposed end-to-end convolutional neural network (CNN) based model extracts contextual information by combining low-level and high-level features. We trained the proposed model using huge publicly available databases, (CBIS-DDSM, BCDR-01, and INbreast), and a private database from the University of Connecticut Health Center (UCHC).

**Results:** We compared the performance of the proposed model with those of the state-of-the-art DL models including the fully convolutional network (FCN), SegNet, Dilated-Net, original U-Net, and Faster R-CNN models and the conventional region growing (RG) method. The proposed Vanilla U-Net model outperforms the Faster R-CNN model significantly in terms of the runtime and the Intersection over Union metric (IOU). Training with digitized film-based and fully digitized MG images, the proposed Vanilla U-Net model achieves a mean test accuracy of 92.6%. The proposed model achieves a mean Dice coefficient index (DI) of 0.951 and a mean IOU of 0.909 that show how close the output segments are to the corresponding lesions in the ground truth maps. Data augmentation has been very effective in our experiments resulting in an increase in the mean DI and the mean IOU from 0.922 to 0.951 and 0.856 to 0.909, respectively.

**Conclusions:** The proposed Vanilla U-Net based model can be used for precise segmentation of masses in MG images. This is because the segmentation process incorporates more multi-scale spatial context, and captures more local and global context to predict a precise pixel-wise segmentation map of an input full MG image. These detected maps can help radiologists in differentiating benign and malignant lesions depend on the lesion shapes. We show that using transfer learning, introducing augmentation, and modifying the architecture of the original model results in better performance in terms of the mean accuracy, the mean DI, and the mean IOU in detecting mass lesion compared to the other DL and the conventional models.

## Background

Breast cancer is the second most common cause of cancer death among women in the United States [1]. According to the American cancer society, the female breast cancer death rate declined by 38% from its maximum in 1989 to 2014 (avoiding about 300,000 deaths) [1]. In 2012, the estimated number of deaths among females in the USA is 43,909 out of 293,353 of all cancer deaths. Moreover, in 2017, it is estimated that there will be 40,610 breast cancer deaths in the USA [1, 2]. This decline in mortality is partially due to the advances in mammography screening and conventional computer-aided diagnosis models (CAD) [3, 4]. In the last few years, deep learning (DL) models and, in particular, convolutional neural networks (CNNs) have achieved state-of-the-art performance for image classification, lesion detection for mammography [5–7], and for medical applications in general [8]. Various approaches have been proposed to further improve the accuracy of deep CNNs [6, 7].

In a recent survey [9] on conventional CAD models and DL classification models for mammograms (MGs) images, it has been shown that conventional models have limitations in classifying MG images. Recent research studies in [4, 10–12] present different conventional models to detect lesions in MG images. Most of the conventional models depend on a pre-requisite set of local hand-crafted features that cannot be generalized to work on a new data-set. Conventional CAD models consider limited feature types (e.g. texture features, shape features, and grey level intensity features), which require expert knowledge for selecting them [4,9,11,12]. Poor feature extraction and selection cause challenge to build a successful classifier [4, 6, 7, 9–12]. However, the state-of-the-art CNNs, extract global features from MG images [6,7,13]. In CNNs, the first layers of the network capture basic coarse features such as oriented edges, corners, textures, and lines while subsequent layers construct complex structures or global features [5].

Despite the initial success of DL models for the segmentation of lesions in medical images as general, the segmentation of lesions in mammography using DL methods has not been studied thoroughly. A few studies have used a CNN-based model for lesion segmentation [14, 15] in MGs and more research need to be done in this topic [6–8]. Few studies have employed CNN-based models for lesion detection and localization [14, 16–28]. These detectors provide bounding boxes (BBs) indicating regions of interests (ROIs), not real lesion segments. The region-based CNN (R-CNN) models [29] and its faster variants, Fast R-CNN [30], and Faster R-CNN [31] have recently become more popular for localization tasks in mammography [18–22]. Although these detectors offer compelling advantages, training R-CNN is time-consuming and memory expensive. In R-CNN [29], the whole process involves training three independent models separately without much-shared computation: 1-the CNN for feature extraction, 2-the top SVM classifier for identifying ROIs’ and 3-the regression model for tightening region BBs. The R-CNN [29] uses the Selective Search method [32] to first generate initial sub-segmentations and generate candidate regions, then it uses the greedy algorithm to recursively combine similar regions into larger ones, and lastly uses the generated regions to produce the final candidate region proposals. These region proposals lower down the number of the potential BBs [18, 19].

Instead of extracting CNN feature vectors independently for each region proposal, the Fast R-CNN [30] aggregates them into one CNN forward pass over the entire image and the region proposals share this feature matrix. Then the same feature matrix is used for learning the object classifier and the BB regressor. In R-CNN and Fast R-CNN, the region proposals are created using the Selective Search method, which is a slow process that is found to be the bottleneck of the overall detection and the localization process. The Faster R-CNN [31] is a better approach that constructs a single unified model composed of region proposal network (RPN) and Fast R-CNN with shared convolutional feature layers. The RPN is a fully convolutional network (FCN) that is trained to generate region proposals, which are then used by the Fast R-CNN for detection. The time cost of generating region proposals is much smaller in the case of RPN than Selective Search, as RPN shares the most computation with the object detection network using the shared convolution layers [30, 31].

The mask R-CNN for simultaneously detecting and segmenting object instances in an image is proposed in [33]. This model extends the Faster R-CNN model by adding a branch which is a FCN for predicting an object mask in parallel with the existing branch for BB recognition. A mass detector has been refined using a cascade of R-CNN and RF classifiers and an additional stage to eliminate false positives [21].

Patch-based CNNs [16,17,23,34] were also proposed to detect masses. In [16], every breast image is divided into patches, and each patch is tested with the CNN model individually. The final detection of lesions in each case is based on the overall scores of all the patches. In [25–27] the famous YOLO CNN (You Only Look Once) [35] is used for breast mass classification and localization. YOLO [35] is a single end-to-end CNN that predicts BBs and class probabilities directly from full images in one evaluation.

Recently, the FCN and its variant improved models as U-Net [36], SegNet [37], Dilated-Net [38], have yielded outstanding results for semantic segmentation of bio-medical images and natural images [6, 13, 14]. These semantic segmentation networks are based on encoding (convolutional) and decoding (de-convolutional) layers. These approaches avoid using the fully connected layers (FCLs) of CNNs to convert the image classification networks into image semantic segmentation networks.

In this study, we developed a new model based on the architecture of the semantic segmentation U-Net model [36] to precisely segment mass lesions in MG images. In the proposed architecture, we used a pre-trained encoder layers and we added batch normalization layers (BN) [39], and dropout layers [40]. U-Net [36] is an end-to-end model that takes an image, find automated features in each layer, detects, and segments breast lesion using a single model and a unified training process. We trained the proposed Vanilla U-Net model using large public data-sets (CBIS-DDSM [41], BCDR-01 [42], and INbreast [43]). We applied data augmentation (Aug.) to the training images to present the lesions in many different sizes, positions, angles. To enhance the contrast of the MGs, we applied image pre-processing before training the proposed model. We compared the performance of the proposed segmentation model in detecting lesions with those of the state-of-the-art Faster R-CNN [18], the conventional region growing (RG) [44], FCN [45], Dilated-Net [38], original U-Net [36], and SegNet [37] models.

## Material and methods

### Databases

We conducted our experiments on four databases, CBIS-DDSM [41], INbreast [43], UCHCDM [46], and the BCDR-01 [42]. CBIS-DDSM [41] is a digitized screen-film mammography (SFM) database that is a subset of the digitized DDSM database [47] with updated lesion segmentation and BBs, and verified pathology. We used 1,696 images from the CBIS-DDSM database that have mass lesions. BCDR-D01 is an SFM repository with 64 patients and 246 MGs [42]. In total, we used 136 mass segmentation from this database to conduct our experiments. The INbreast is another public database for MGs which comprises fully field digital mammography (FFDM) images [43]. It has a total of 410 images, and we used 116 MGs that are annotated for masses. UCHCDM is a private database of FFDM images collected from the University of Connecticut health center (UCHC) [46, 48]. In total, the UCHCDM database consists of 173 patients with 1,340 FFDM images. We selected 59 cases out of the 173 that have mass lesions, with a total of 118 MGs with mass annotations. The CBIS-DDSM, IN-breast, and UCHCDM datasets include separate files that show region of interest (ROI) annotations for the abnormalities, provided by radiologists [6, 7].

We combined these databases and generated a new dataset containing MGs with different resolutions (see supplementary Fig.1, Additional file 1). This new data-set provides mass lesions of different sizes, shapes, and margins. All images containing suspicious areas have associated pixel-level ground truth maps (GTMs) indicating the true locations of suspicious regions (see supplementary Fig.2, Additional file 1). The total number of images used in this combined dataset is 2,066 and each image has its corresponding GTM. We divided the images into a training data-set of 1,714 images, validation data-set of 204 images, and test data-set of 148 images. Images reserved for testing were not used in the training and the validation data-set. Images that come from the same patient were not split across the training and test data-sets.

### Pre-processing

Pre-processing of MGs is an essential step before applying DL methods. Its main goal is to enhance the characteristics of MGs by applying a set of filters to improve the performance of the downstream analysis. First, we detect the breast boundary for removing a big portion of the black background [49,50]. After that, we apply the adaptive median filter (AMF) [51] to remove any existing noises. Then, we employ the contrast limited adaptive histogram equalization (CLAHE) [52] to enhance the contrast of the MGs [6, 49, 50], see supplementary Pre-processing subsection, Additional file 1. The superior performance of the CLAHE filter compared to other filters are shown in [6, 53]. All full MGs are converted into png format and re-sized to 512*×*512.

### Data augmentation

In this study, we adopted augmentation techniques to increase the size of our training data-set to avoid overfitting the model. We adopted the augmentation techniques used in [54–56]. We generated augmented images by image rotation in a range of ± 10 degrees, left-right flips, translate images left and right by 10%, translate images up and down by 10%, and zoom in and out by 20%. The mass segmentation maps are represented by binary images that are cropped, re-sized and augmented in the same way as their corresponding MGs. All pixels in the GTMs are labeled as belonging to background or breast lesion classes. The size of the generated augmented data-set is ten times larger than the size of the original data-set.

### Semantic segmentation using U-Net

The U-Net is a popular end-to-end encoder-decoder network for semantic segmentation that is originally invented for bio-medical image segmentation tasks [36]. U-Net [36] extends the FCN [45] with a U-shape architecture, which allows features from shallower layers to combine with those from deeper layers. U-Net consists of a contracting path to capture features and an asymmetric expanding path that enables precise localization and segmentation of pixels. This architecture has a U shaped skipping connections that connect the high-resolution features from the contracting path to the up-sampled outputs of expanding path. After collecting the required features in the encoding path, the decoding path performs nonlinear up-sampling of the feature maps before merging with the skip connections from the encoding path followed by two 3×3 convolutions, each followed by an element-wise rectified linear unit (ReLU). The skip concatenation allows the decoder at each stage to learn back relevant features that are lost when pooled in the encoder. The final output is obtained by passing the result through a pixel-wise Softmax classifier after the last convolution layer, which independently assigns a probability to each pixel.

### Architectural modifications

We have modified the original U-Net model [36] to improve its performance for the task of segmenting lesions. We added BN layers [39], dropout layers [40], and increased the number of convolution layers. We also trained the proposed model with augmented data-set. In our implementation, we used a pre-trained VGG-16 model [57] on ImageNet as the encoder portion of the proposed Vanilla U-Net model and thus can benefit from the features created in the encoder. Studies have shown that transfer learning techniques from one domain to another are very effective to boost the performance of the current task [6,7]. VGG-16 [57] consists of seven convolutional layers, each followed by a ReLU activation function, and five max-polling operations. The first convolutional layer of the VGG-16 model produces 64 channels and then, as the network deepens, the number of channels doubles after each max pooling operation until it reaches 512. On the following layers, the number of channels does not change. To construct the encoder part of the Vanilla U-Net, we removed the last FCLs of the VGG-16 model and replace them with two convolutional layers of 512 channels that serves as a bottleneck part of the network, connecting the encoder with the decoder.

Figure 1 shows our modified model. The encoding path consists of five convolutional layers which perform convolution with a filter bank to produce a set of feature maps. A BN layer is added between the convolution layer and the ReLU layer. Batch normalization [39] prevents internal covariate shifts as data are filtered through the network, and it reduces the training time, prevents data overfitting, helps stack more layers, and generally increases the performance of deep CNNs. We added drop-out layers of 0.5 after each convolutional layer to help regularize the networks [40]. Following that, max-pooling with a 2×2 window and stride 2 is performed and the resulting output is sub-sampled by a factor of 2. The max-pooling layer reduces the dimensionality of the resulting output, enabling the further collection of features. To construct the decoder, we used transposed convolutions layers that doubles the size of the feature maps while reducing the number of channels by half. The output of a transposed convolution at each level is then concatenated with an output of the corresponding part of the decoder at the same level. Also, to keep the size of the output map the same as the size of the original input MGs, a padded convolution is applied to keep the dimensions consistent across concatenation levels.

**Fig. 1.**
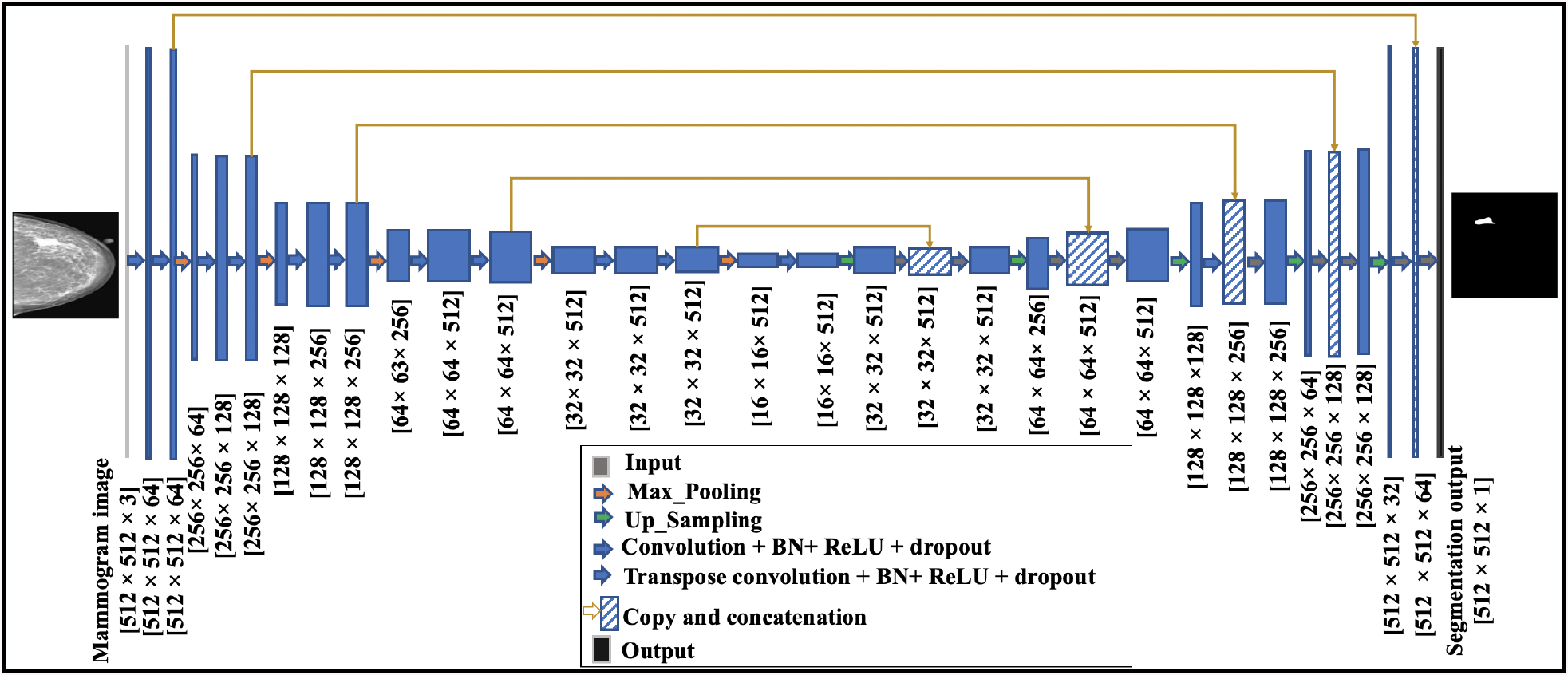
The U-Net architecture consists of convolutional encoding and decoding units that take an image as input and produce the segmentation feature maps with respective pixel classes. The yellow arrows show the skip connections between the layers.

Our data-set has imbalanced data representation. In an imbalanced representation, classes are represented by significantly different numbers of pixels, which makes the learning algorithm biased towards the dominating class (i.e. breast tissues and/or background). We address this problem by introducing class weights into the Dice loss function [58]. The class weight is the ratio of the median of class frequencies computed on the entire training set divided by the class frequency [58]. This implies that the breast tissues and background class in the training set have weights smaller than the weights of the lesion class. Moreover, we applied the augmentation techniques explained in the previous sub-section, instead of applying elastic deformations as done in the original U-Net model [36].

For training, the Dice loss function was minimized using Adam optimizer [59] with a decreasing learning rate (LR) initialized to 1*e^−^*^2^ and a momentum of 0.9. We used the famous early stopping technique to avoid over-fitting the model by monitoring the DI value of the validation data-set. The training of the models stops when DI is not improved every 20 epochs. Before each epoch, the training set is shuffled and every 4 mini-batch images are then picked thus ensuring that each image is used only once in an epoch. We used input MGs resized to 512×512. We developed, trained, and tested the DL models using MATLAB version 2019b. Training and testing the models were done on a Tesla K40m Nvidia graphics processing unit.

### Evaluation metrics

To evaluate the performance of the DL models, the Dice index coefficient (DI), also known as the F1 score, and the Intersection over Union (IOU), also known as the Jaccard index, metrics are used to compare the automated predicted maps with the GTMs [60–62]. We mapped the class probabilities from the Softmax output to discrete class labels and used it to calculate the commonly used DI and IOU metrics, Eqs. 1 and 2, respectively.

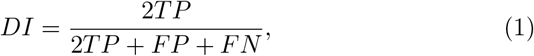

where TP is the number of true positive pixels, FP is the number of false positives and FN is the number of false negatives.

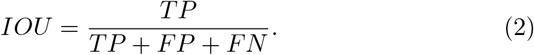

Dice index measures the similarity between the segmented lesions, that have irregular boundaries, and the annotated ground truth maps. IOU measures the intersection ratio between the obtained segmentation BBs and the ground truths BBs. Thus, IOU is used for localization of lesions and is best with rectangular boundaries. The output of different segmentation models might have similar IOU (lesion well localized) but with slightly different DI value that show how precise the lesions within the MG image are segmented.

The DI score gives more weight to TPs than FPs and FNs (Eq. 1). While IOU score gives more weight to TPs, FPs, and FNs (Eq. 2). Similar to DI, the IOU score ranges from 0: 1, with 0 signifying no overlap and 1 signifying perfectly overlapping segmentation [62]. Also, for each class, IOU can be calculated using the ratio of correctly classified pixels to the total number of ground truth and predicted pixels in that class (Eq. 2). The mean IOU of each class is weighted by the number of pixels in that class.

As mentioned in the Background section, most of the lesion detection models provide BBs for an indication of a region with an abnormality. To compare the performance of the proposed Vanilla U-Net model with detection models providing BBs such as the Faster R-CNN, a BB is generated around every detected lesion. The BBs are generated based on a minimum and maximum points of x and y coordinates, which indicate the locations of masses. We calculated the accuracy of localization by considering the detected segment and BB as TP if the center of the segment or the BB overlaps with the ground truth by more than 50%. For each class, the accuracy metric is the ratio of correctly classified pixels to the total number of pixels in that class, according to the GTMs (Eq. 3). Mean accuracy is the average accuracy of all classes in all images.

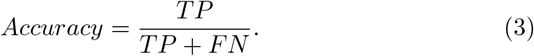

We also calculated the Boundary F1 contour matching score (BF-score) for each image, which indicates how well the predicted boundary of each class aligns with the true boundary. For each class, the mean BF-score shows the average BF-score of all classes in all images. Values near 1 means perfect boundary.

## Results

### Comparison with state-of-the-art methods

We adopted the Faster R-CNN model [18], original U-Net [36], VGG16-based FCN-8s model [45], VGG16-based SegNet model [37], Dilated-Net [38] model, and the conventional RG CAD model [4, 44] to apply to MGs for comparing their performances with that of our model in terms of mean accuracy, mean DI, mean IOU, mean BF-score, and the inference time in seconds per image (see Table 1). The architecture of these models in more detail is given in the Additional file 1. We trained the Faster R-CNN detector proposed in [18] to detect breast cancer lesions on MGs using our augmented data-set. We also implemented the RG method proposed in [44] and apply it to our MG images.

**Table 1.** The performance of the proposed Vanilla U-Net model, original U-Net, Faster R-CNN, SegNet, Dilated-Net, FCN, and RG.

| Model | Mean accuracy | Mean DI | Mean IOU | Mean BFscore | Mean inference time (second)/image |
| --- | --- | --- | --- | --- | --- |
| -Proposed Vanilla U-Net, with Aug. | 0.926 | 0.951 | 0.909 | 0.964 | 0.115 |
| -Proposed Vanilla U-Net, without Aug. | 0.910 | 0.922 | 0.856 | 0.940 | 0.115 |
| -Original U-Net, with Aug. | 0.842 | 0.818 | 0.693 | 0.800 | 0.118 |
| -Original U-Net, without Aug. | 0.821 | 0.801 | 0.668 | 0.776 | 0.118 |
| -Faster R-CNN, with Aug. | 0.702 | - | 0.601 | - | 0.454 |
| -SegNet, with Aug. | 0.853 | 0.824 | 0.701 | 0.822 | 0.098 |
| -Dilated-Net, with Aug. | 0.832 | 0.799 | 0.665 | 0.701 | 0.094 |
| -FCN, with Aug. | 0.843 | 0.802 | 0.669 | 0.752 | 0.091 |
| -RG | 0.801 | 0.602 | 0.401 | 0.603 | 0.320 |

The test data-set consists of SFM and FFDM MG images. Figures 3 (b) and 4 (b) show the original FFDM MG images from the INbreast database. Figure 2 (b) shows the SFM MG images from the DDSM database. Where the red BBs in Figs. 2 (b), 3 (b) and 4 (b) show the ground truth given by radiologists. The calculated DI and/or IOU for each detection is shown under each image.

**Fig. 2.**
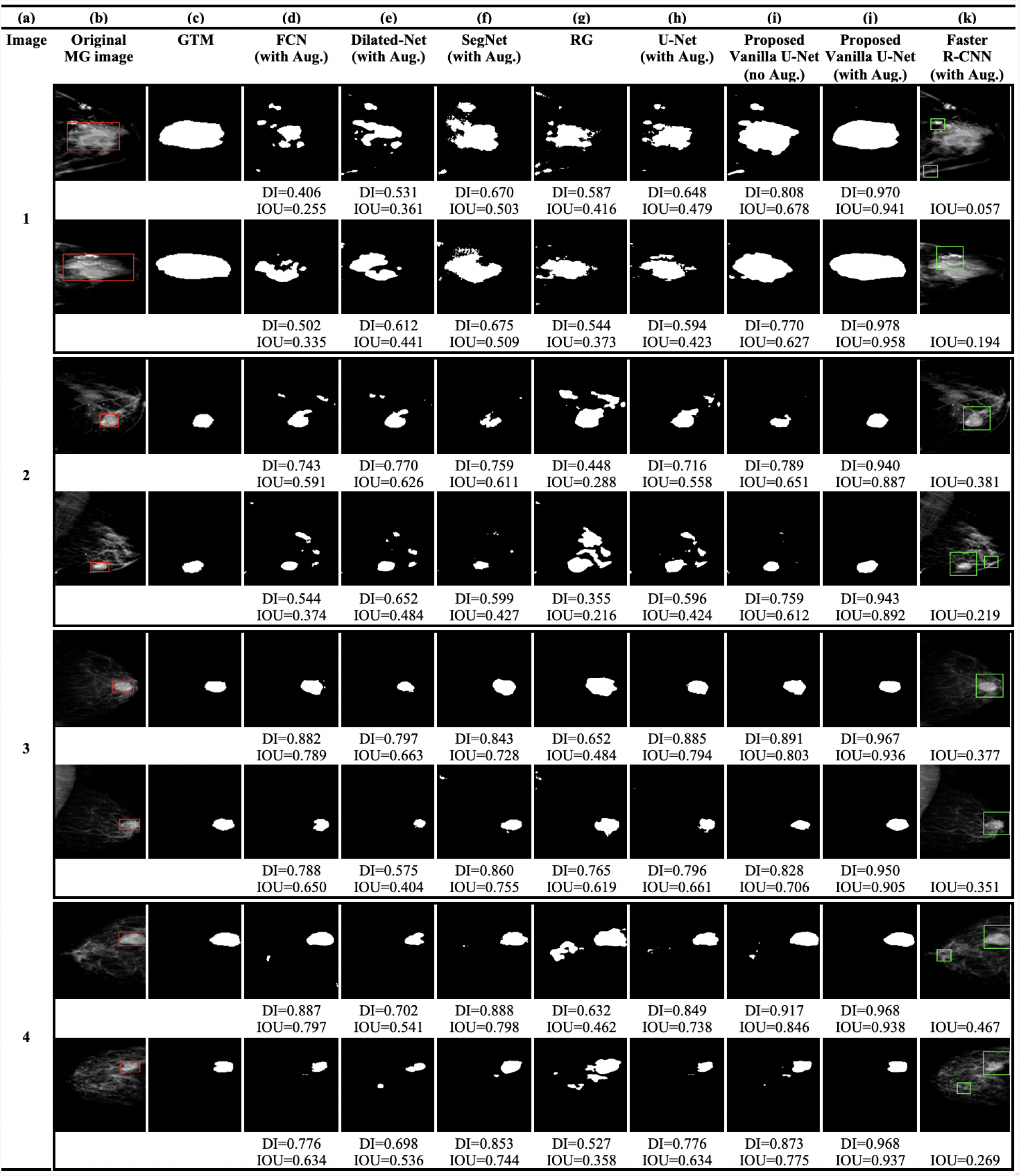
(a) Image index, (b) the original SFM MG images from the DDSM database (the red rectangles show the location or the BBs of the ground truth lesions), (c) the GTMs given by radiologists, (d) the FCN model, (e) the Dilated-Net model, (f) the SegNet model, (g) the RG method, (h) U-Net model trained with the augmented data-set, (i) the proposed Vanilla U-Net model without augmentation, (j) the proposed Vanilla U-Net model trained with the augmented data-set, and finally (k) the Faster R-CNN model trained with the augmented data-set.

**Fig. 3.**
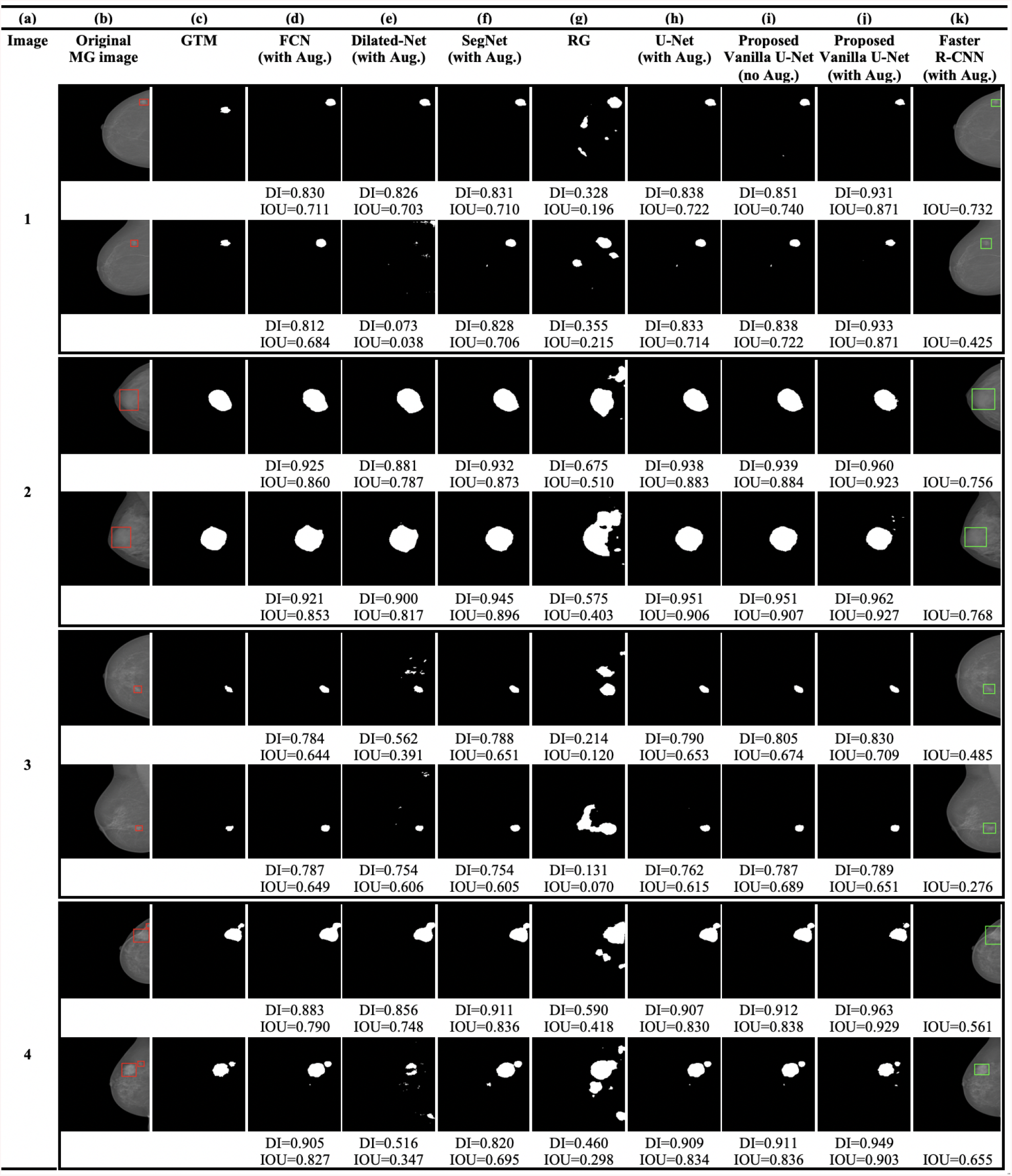
(a) Image index, (b) the original FFDM MG images from the INbreast database (the red rectangles show the location or the BBs of the ground truth lesions), (c) the GTMs given by radiologists, (d) the FCN model, (e) the Dilated-Net model, (f) the SegNet model, (g) the RG method, (h) U-Net model trained with the augmented data-set, (i) the proposed Vanilla U-Net model without augmentation, (j) the proposed Vanilla U-Net model trained with the augmented data-set, and finally (k) the Faster R-CNN model trained with the augmented data-set.

Table 1 shows the evaluation metrics of all the networks included in this study in terms of mean accuracy, mean DI, mean IOU, mean, mean BF-score, and mean inference time (second)/image. In Table 1, the performance of the models is shown for the detected segments/tight BBs in comparison with the GTMs. The mean DI and the mean IOU of the proposed Vanilla U-Net are 0.951 and 0.909, respectively, which are higher compared to other models (Table 1). The BF-score of the proposed Vanilla U-Net model is 0.964 which exceeds the other segmentation models.

The architecture of the SegNet model is much closer to that of the U-Net model compared to the other segmentation models. However, the boundary of the detected regions of SegNet model is not aligned with the true boundary (Figs. 2 (f), 3 (f) and 4 (f)). The Seg-Net model has a BF-score of 0.822. The SegNet model performs better when detecting lesions in FFDM MGs compared to SFM MGs. In contrast, the proposed Vanilla U-Net model performs very well for both kinds of images (Figs. 2 (j), 3 (j) and 4 (j)). The proposed Vanilla U-Net shows better performance compared to SegNet. U-Net transfers the entire feature maps to the corresponding decoders and concatenates them to the up-sampled decoder feature maps, which gives precise segmentation. SegNet has much fewer trainable parameters compared to the U-Net model since the decoder layers use max-pooling indices from corresponding encoder layers to perform sparse upsampling. This reduces the inference time at the decoder expanding path since the generated encoder feature maps are not involved in the upsampling. Thus, the SegNet model reveals a trade-off between the memory versus accuracy involved in achieving good segmentation performance (Table 1). The mean DI and the IOU of the trained SegNet model on augmented data-set are 0.824 and 0.701, respectively, compared to 0.952 and 0.909 of the U-Net model.

The trained Dilated-Net has a mean DI of 0.799, a mean IOU of 0.665, respectively. Moreover, its BF-score is 0.701 that is lower than that of the proposed Vanilla U-Net model and the SegNet model BF-score. Also, the performance of the Dilated-Net model is worse in the case of SFM images (Fig. 2 (e)). Even-though some images in Figs. 2 (e), 3 (e) and 4 (e) show slightly better DI than that of SegNet, the performance of the model on all the test data-set is lower than that of the SegNet model. In contrast to U-Net and SegNet, down-sampling layers are not required in the Dilated-Net to obtain large receptive fields and hence, high-resolution maps can be directly predicted by the model. Down-sampling layers are widely used for maintaining invariance and controlling overfitting of the model, however it reduces the spatial resolution. To retrieve the lost spatial information, the Up-sampling layers in U-Net and SegNet are used, but with additional memory and time constraints.

We also adapted the FCN-8s VGG16 based network [45] to compare its performance with that of the proposed Vanilla U-Net model. FCN-8s up-samples the final feature map by a factor of 8 after fusing feature maps from the third and fourth max-pooling layers, thus having better segmentation than its variants. The FCN in our study has a mean DI of 0.802 and a mean IOU of 0.669, respectively. Moreover, the BF-score of the best trained FCN is 0.752 which is lower than that of the proposed Vanilla U-Net by 0.212. The mean DI scores of the Dilated-Net and the FCN model are close for some of the images, however, the FCN give the lowest scores among all the segmentation DL models.

As we mentioned in the Method section, we generated tight BBs surrounding detected segments to compare the performance of the proposed model with that the BB-based models such as Faster R-CNN. The proposed Vanilla U-Net model shows better performance in detecting true segments compared to the Faster R-CNN model as shown in Fig. 2 (i: k), Fig. 3 (i: k) and Fig. 4 (i: k). In Fig. 2, the Faster R-CNN model introduces FPs in the SFM images as in rows (1, 2, and 4 (k)). In Fig. 4 (k), the Faster R-CNN model introduces some FPs as in row (2k and 4k), as an example. The proposed Vanilla U-Net model shows better performance with both FFDM and SFM images than the Faster R-CNN model. To have a better understanding of the performance of the proposed Vanilla U-Net and other models, we included the DI and/or the IOU under every image. We also show the detection of the proposed Vanilla U-Net for every CC and MLO view of the same patient. The IOU of the proposed model exceeds the IOU of the Faster R-CNN by 0.308, as shown in Table 1. We considered the detected BBs as TP if the center of the detected BB overlaps with the ground truth BB with greater than 50%.

**Fig. 4.**
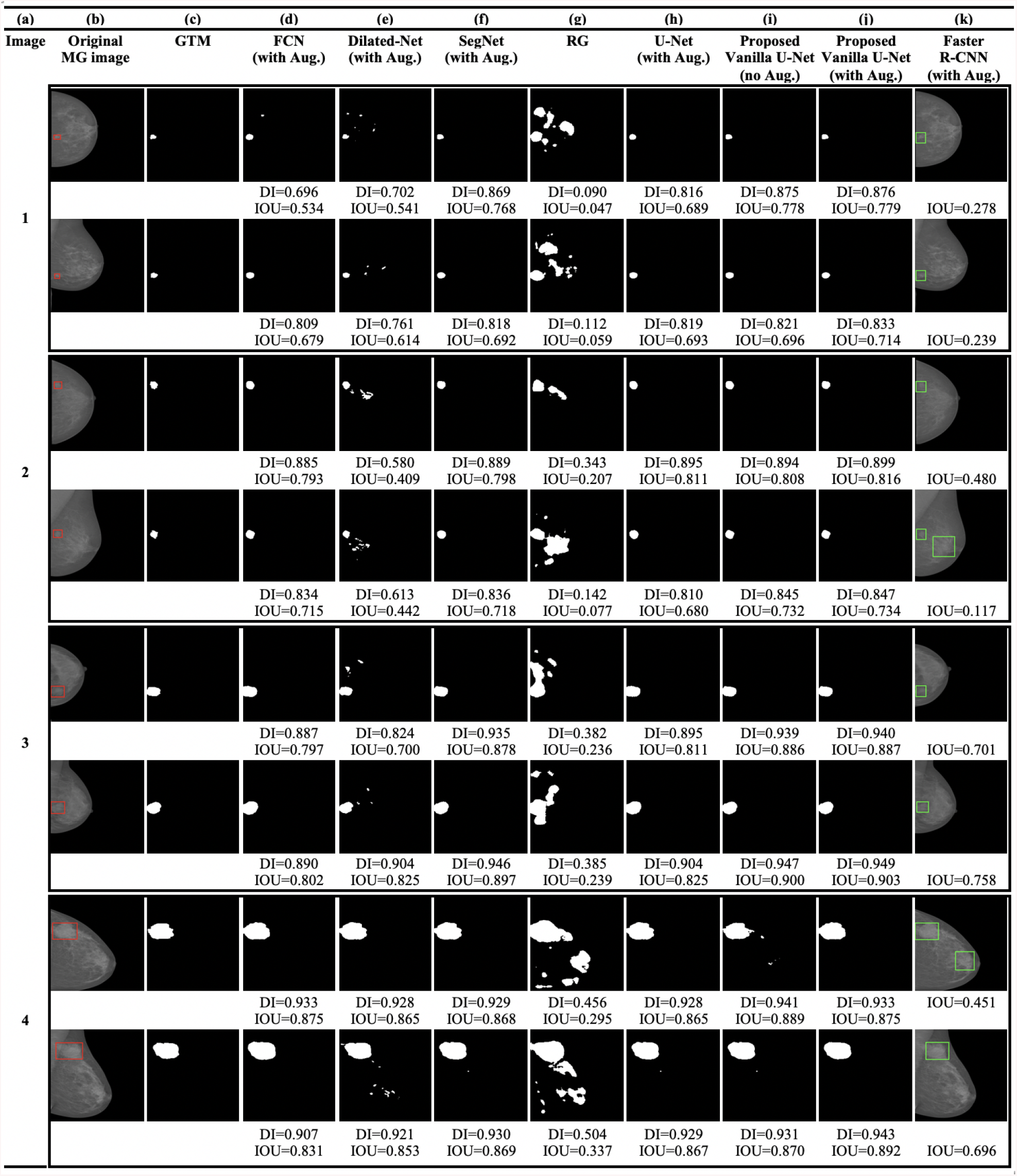
(a) Image index, (b) the original FFDM MG images from the INbreast database (the red rectangles show the location or the BBs of the ground truth lesions), (c) the GTMs given by radiologists, (d) the FCN model, (e) the Dilated-Net model, (f) the SegNet model, (g) the RG method, (h) U-Net model trained with the augmented data-set, (i) the proposed Vanilla U-Net model without augmentation, (j) the proposed Vanilla U-Net model trained with the augmented data-set, and finally (k) the Faster R-CNN model trained with the augmented data-set.

The accurate automated seed selection process is very important for lesion segmentation. As RG segmentation’s results are sensitive to the initial seed pixels, the final segmentation results would be incorrect if the seeds are not properly selected by the automated process. The RG method works better when it is used with patches of images that contain the ROI because the initial seed pixels are close to the center of the ROI. Figures 2 (g), 3 (g) and 4 (g) show the detection using the RG method. Figures 2 (g), 3 (g) and 4 (g) show that the DL models outperform the conventional CAD models in terms of DI in the segmentation of tumors in whole images. The mean DI and mean IOU of the RG method are 0.602 and 0.401, respectively.

We also explored the current state-of-the-art DL models for segmentation or localization of lesions in MG images through a literature survey [6]. The reported performance metrics of several models are shown in Supplementary Table 1, Additional file 1. The researchers used various metrics to report their work, which makes a direct comparison between these different approaches difficult. Moreover, the number of training data-set and the size of training images vary from a study to another one. However, it gives us some insights into the strategies used in these studies. Researchers who applied the transfer learning (TL) strategy to train their DL models reported that the TL approach helped them to report better detection accuracy, see supplementary Table 1, Additional file 1. Moreover, the size of the training dateset and the resolution of the MGs play an important role in increasing the model’ accuracy [63].

### Effect of augmentation

In our experiments, we observed that the mean DI of the proposed Vanilla U-Net model increased slightly from that of the original U-Net model when we added BN layers or used dropout layers or increased the number of convolution layers, one at a time. Moreover, we observed that the proposed modifications, together, have increased the mean DI of the proposed Vanilla U-Net model in comparison with that of the original U-Net model from 0.801 to 0.951. But mostly the augmentation of the data-set had a great impact on the performance of the proposed Vanilla model in terms of mean DI. And because of that, we investigated the effect of augmentation in the performance of the proposed Vanilla U-Net model.

The augmented training data-set results in 17,140 images. Figures 2 (i: j), 3 (i: j), and 4 (i: j), illustrate the effect of augmentation on the proposed Vanilla U-Net model. For example, the values of the DI of the augmented model, as shown in (j), are higher than the ones of the trained model without augmentation, as shown in (i). Table 2 shows the improvement in terms of DI for both training and validation data-sets when using augmented data-set compared to when using the original one. The DI improves from 0.910 (training), and 0.842 (validation) to 0.972 (training), and 0.942 (validation). The augmented data-set also affect the localization precision significantly (Table 1). The BF-score improves from 0.940 to 0.964 in the case of the proposed augmented U-Net model.

**Table 2.** The performance of the proposed Vanilla U-Net model.

| Model | Training |  | Validation |  | Test Dice |
| --- | --- | --- | --- | --- | --- |
|  | Dice | Loss | Dice | Loss |  |
| -Proposed Vanilla U-Net, with Aug. | 0.972 | 0.016 | 0.942 | 0.024 | 0.951 |
| -Proposed Vanilla U-Net, no Aug. | 0.910 | 0.064 | 0.842 | 0.144 | 0.922 |

Figure 5 shows that the histogram of the mean of IOU value for the test images increases using the proposed Vanilla U-Net model after data augmentation. The mean of IOUs of the proposed Vanilla U-Net improves from 0.856 to 0.909 when training with the augmented data-set (Fig. 5 and Table 1). In general, the performance of the DL techniques improves as the size of the training data-set increases [6, 7]. Figures 2 (i: j), 3 (i: j), and 4 (i: j), show that the DI per image increases when the proposed model is trained with the augmented mixed data-set. The FP pixels decreased in the case of the augmented model, as shown in rows (1, 2, and 4) in Fig. 2 (i: j).

**Fig. 5.**
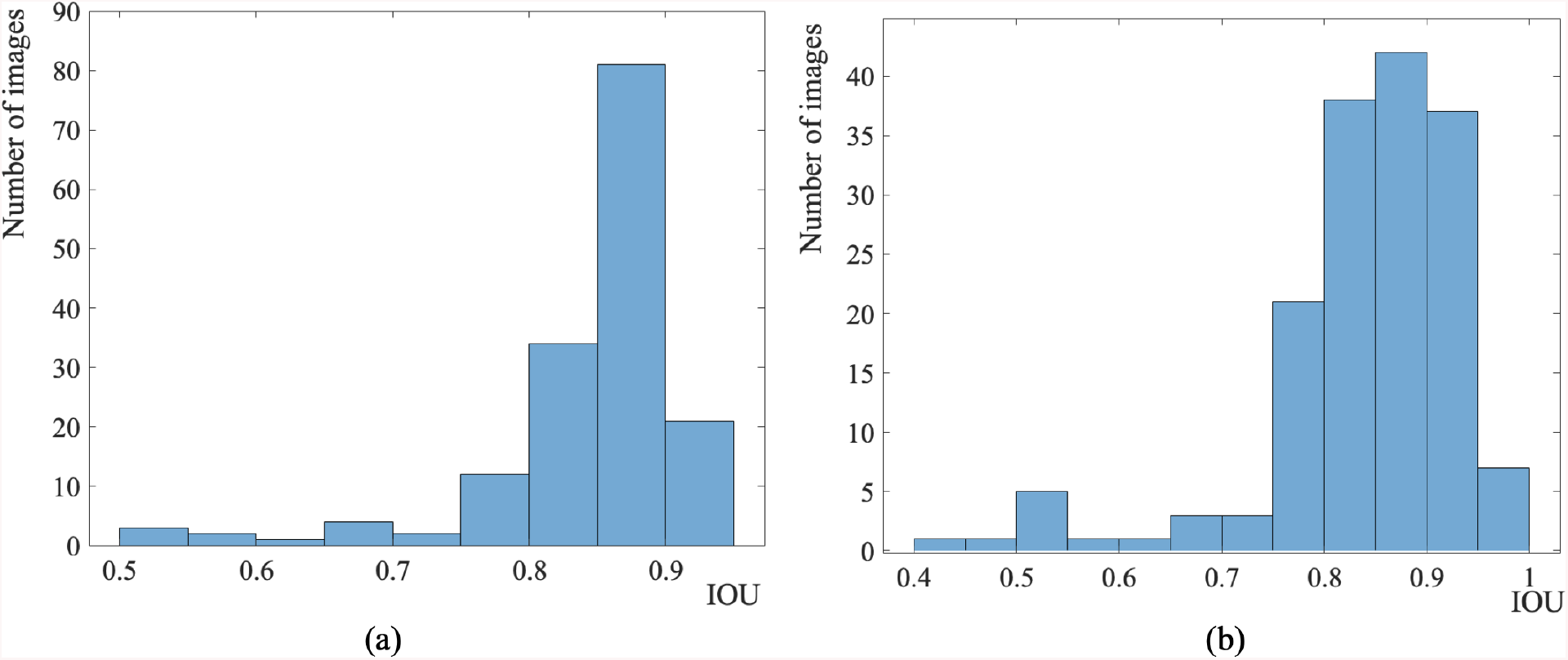
Histogram of the mean of IOU value for the test images using the proposed Vanilla U-Net model before Aug. (a) and after Aug. (b).

### Effect of image size and data-set size

One of the factors that make a localization model or a semantic segmentation model superior to other models, is its ability to help the radiologists to detect small lesions that can be missed with the naked eye. A recent study in [63] on MGs shows that the resolution of the training images affects the performance of the CNN model. Recent studies, as shown in Table 1, Additional file 1, use MGs of small sizes as 40×40 and 227×227. The standard image sizes of 224×224, and 227×227 are used excessively for training CNNs to detect objects in natural images [61]. However, the requirement to find small mass lesions in aggressively down-sampled high-resolution images is unlikely to be successful for MGs [6, 63].

In our initial work in [14], we trained the proposed model with images of size 256×256 and found that the proposed model failed to find small lesions in images of high density. As a result, we changed our training strategy to include MGs of size 512×512 instead of 256×256. Figure 6 shows some FFDM test images that have small lesions that are detected with DI greater than 50%.

**Fig. 6.**
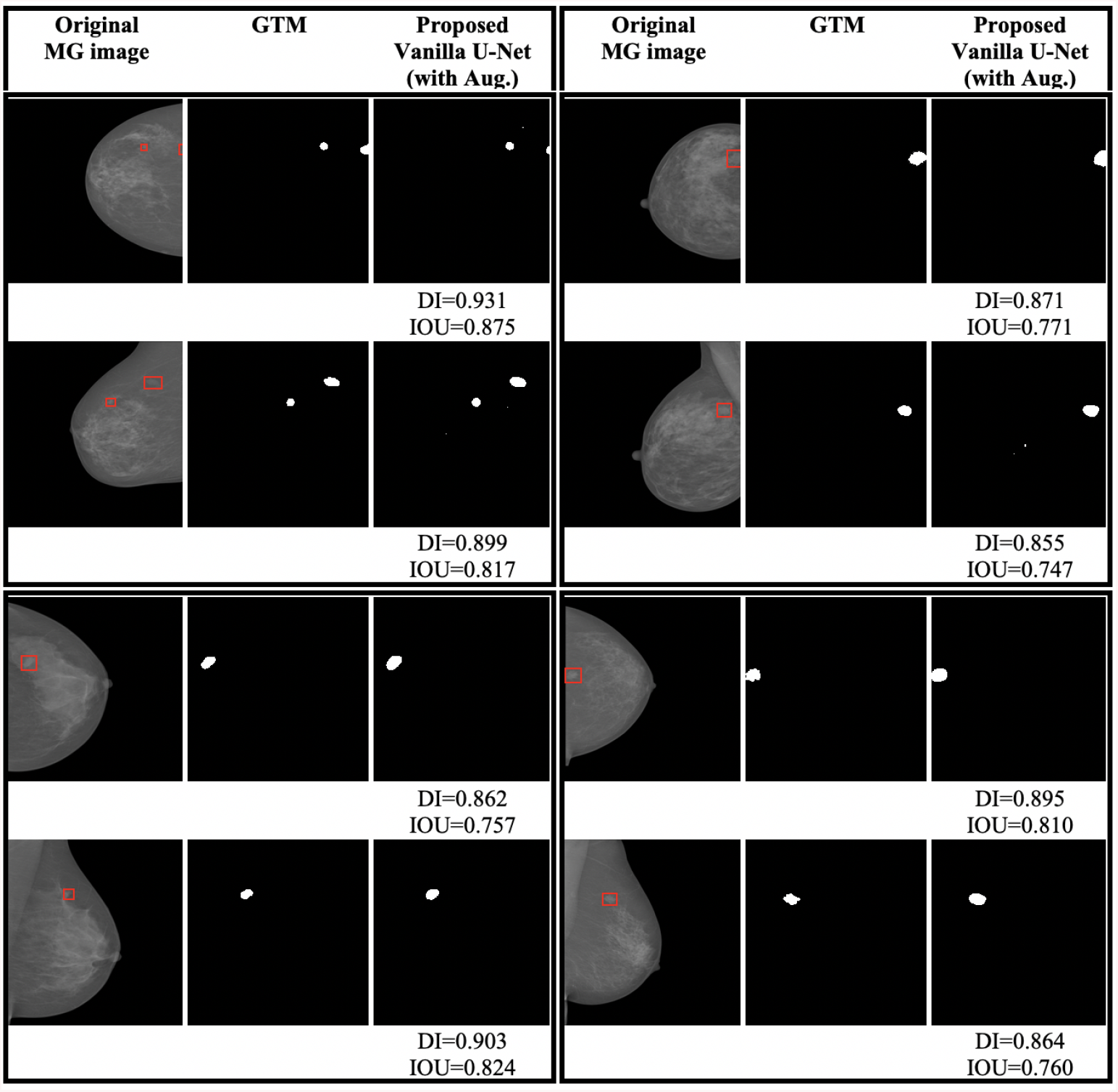
Test cases with small lesions.

Because of the architecture of the proposed Vanilla model, images with a side divisible by 32 (e.g. 1024×1024) can be used as an input to the current network implementation. In the future, we will conduct our experiments on high-resolution images to get a competitive performance to recent state-of-the-art models as the size of MG images in the clinical settings are generally larger than 1024×1024 [7].

### Improvements of the proposed model over the original U-Net model

The proposed model yields an improvement of 16.32% in the mean DI and 31.16% in the mean IOU, respectively, relative to that of the original U-Net model (Table 1). The original U-Net model is trained from scratch. Moreover, increasing the data-set size by using the proposed augmentation technique improves the segmentation’s quality (BF-score yields an increase of 20.5% relative to that of the original U-Net model). The original U-Net did not use the BN technique. Original U-Net did not use the BN technique. Batch normalization helps the proposed model avoiding vanishing gradient problem, stacking more layers, accelerating training, and using less number of epochs. In the proposed model, we went deeper into the number of layers from four to five convolution layers. By increasing the number of convolution layers, the segmentation process incorporates more multi-scale spatial context and captures more local and global context.

### Timing performance

To assess the runtime performance of these models, we measured the mean inference time per image taken by each model to detect lesions in the test data-set, as shown in Table 1. The proposed Vanilla U-Net model is faster by 0.34 seconds than the Faster R-CNN model [18]. The inference time of the SegNet, Dilated-Net, and FCN is less than the proposed Vanilla U-Net by a fraction of second. Even though the inference time of the RG method is of about 0.3 seconds, it introduces a lot of FPs when tested on whole images as shown in Figs. 2 (g), 3 (g), and 4 (g), and the statistics of Table 1. The proposed Vanilla U-Net model is faster than the Faster R-CNN proposed in [19] and [20], the R-CNN proposed in [21] and [22], and the YOLO model proposed in [27], while proving a high DI, see supplementary Table 1, Additional file 1. We have to emphasize that for radiologists, the accuracy of the proposed CAD or DL model in detecting lesions is the most important feature in the mammography analysis, and the inference time is secondary. An inference time of a fraction of second or even several seconds is not as important as the accuracy of the given model.

## Discussion

We tested our proposed model on SFM and FFDM data-sets for the semantic segmentation of mass lesions in MGs. For our future work, we will consider training the proposed Vanilla U-Net model to detect both the micro-calcification and the mass lesions. We will focus on reducing FP pixels by collecting more data-sets and use higher resolution mammogram images. Finally, we want to use the proposed Vanilla U-Net model to distinguish between benign and malignant breast tumors in mammography images by studying the features of the tumors’ segmented regions only.

## Conclusions

We developed a new deep learning (DL) model called Vanilla U-Net, based on the architecture of the semantic segmentation U-Net model to precisely segment mass lesions in mammogram (MG) images. The proposed end-to-end model extracts low-level and high-level features from MG images. The proposed Vanilla U-Net model efficiently predicts a pixel-wise segmentation map of an input full MG due to its modified architecture. We tested our proposed Vanilla U-Net model using film-based and fully-digital MGs. We compared the performance of our proposed model with state-of-the-art DL models namely Faster R-CNN, SegNet, FCN, and Dilated-CNN. We also compared the performance of the proposed model with the conventional region growing method. The proposed Vanilla U-Net model is superior to the segmentation models under study. The proposed Vanilla U-Net model gives a mean intersection over union (IOU) of 0.909 and a mean accuracy of 0.926 while the Faster R-CNN model gives IOU of 0.601 and a mean accuracy of 0.702, respectively. Similar to the Faster R-CNN model, the Vanilla U-Net model is trained on the full MGs. However, the proposed Vanilla U-Net model is faster and runs 0.337 seconds less than the Faster R-CNN model. We show that the proposed model show improvement in the Dice index (DI) and the IOU by 16.3% and 31.16%, respectively, relative to the original model. The proposed models can be further trained to detect micro-calcification in the future. The presented work is a step towards a precise segmentation of mass lesions in mammography. As medical data-sets are increasing and becoming publicly available, future architectures may be trained end-to-end, removing the need for pre-training on non-medical data-sets.

## Abbreviations

DL: deep learning
MG: mammogram
CNNs: convolutional neural networks
CAD: computer-aided detection
ML: machine learning
TL: transfer learning
RG: region growing
SVM: support vector machine
DDSM: digital database for screening mammography
ROIs: region of interests
GTMs: ground truth maps
BCDR: breast cancer digital repository
BN: batch normalization
ReLU: rectified liner unit
SFM: screen-film mammography
FFDM: digital mammography
UCHCDM: university of Connecticut health center digital mammogram
E2E: end-to-end
CLAHE: contrast limited adaptive histogram equalization
AMF: adaptive median filter
R-CNN: region-based convolutional neural network
YOLO: you only look once
RPN: region proposal network
AUC: area under the receiver operating curve
DI: dice index
ACC: accuracy
IOU: intersection over union
TP: true positive
FN: false negative
TN: true negative
FP: false positive
FPR: false positive rate
TPR: true positive rate
Aug: augmentation
FCL: fully connected layer
FCN: fully convolutional network

## Declarations

### Availability of data and materials

The DDSM data-set is available online at http://www.eng.usf.edu/cvprg/Mammography/Database.html.^[41, 47]^

The INbreast data-set can be requested online at http://medicalresearchinescporto.pt/breastresearch/index.php/Get_INbreast_Database.^[43]^ The breast cancer digital repository (BCDR) data-set can be requested online at http://bcdr.inegi.up.pt.^[42]^

### Ethics approval and consent to participate

Not applicable.

### Consent for publication

Not applicable.

### Author’s contributions

DH and SN designed the study. DH implemented the method, performed the experiments, and interpreted the results. DH and SN wrote the manuscript. SN supervised the study. CY provided the UCHCDM data-set. RA, CY, and JB provided feedback and helped shape the research. All authors read and approved the final version of the manuscript.

## Acknowledgements

Not applicable.

## Additional Files

Additional file 1 - Supplementary materials (semantic segmentation using FCN, semantic segmentation using SegNet, semantic segmentation using Dilated-Net, localization using Faster R-CNN, comparison between state-of-the-art DL models, Supplementary Table 1, Supplementary Figures 1-4).

