## Supplemental File for "Convolutional neural network for automated mass segmentation in mammography"

### Supplementary materials

#### Pre-processing

The AMF [1] is a nonlinear filter that removes impulse noise while preserving edges and corners to improve the image quality. The CLAHE filter increases the contrast between the masses and their surrounding tissues [2–5]. The CLAHE [1] filter operates on small regions in the image, called tiles, rather than the entire image. It calculates the contrast for each tile individually producing local histograms. Each tile's contrast is enhanced and the neighboring tiles are then combined using bilinear interpolation to eliminate artificially induced boundaries. The contrast in the homogeneous regions can be limited using a clipLimit factor to avoid amplifying any noise that might be present in the image. We used tiles of [8 8] and a clipLimit factor of 0.005 with the CLAHE technique. Figure 1 shows a sample of the combined data-set we used in our experiments. Figure 2 shows images containing suspicious areas and its associated pixel-level GTMs. All full MGs and GTMs are converted into png format and re-sized to 512×512. All pixels in the GTM are labeled as belonging to the background (0) or breast lesion (255) (see Fig. 2).

#### Semantic segmentation using FCN

Semantic segmentation is an active research area for medical images where deep CNNs are used to classify each pixel in the image individually. Semantic segmentation results in a map image that is segmented by classes. The fully convolutional network (FCN) [6] is an encoder-decoder network. The encoder path uses a pre-trained VGG16 model [7] and transfer their learned representations by fine-tuning to the segmentation task. The decoder path uses up-sampling operations, and replace the final fully connected layer (FCL) with an N×1×1 convolution layer, which output probabilities for N classes. A skip architecture is proposed by [6] where the weights of shallow, fine layer features are combined with deep, coarse layer features to produce accurate and detailed segmentations, as intensive up-sampling can lead to coarse segmentation maps. There are 3 versions of FCN (FCN-32s, FCN-16s, FCN-8s) based on VGG16 network [6]. In this research, we adapt the FCN-8s VGG16 based network [6] to our segmentation task. FCN-8s up-samples the final feature map by a factor of 8 after fusing feature maps from the third and fourth max-pooling layers.

#### Semantic segmentation using SegNet

The SegNet architecture [8] adopts the VGG16 network [7] along with an encoder-decoder framework wherein it drops the FCLs of the network. SegNet shares a similar architecture to the encoder-decoder U-Net described in the previous subsection. However, in SegNet, the indices at each max-pooling layer in the encoder contracting path at each level are stored and later used to up-sample the corresponding feature map in the decoder by unpooling it using those stored indices (Fig. 3). Storing the indices from the contraction path helps keep the high-frequency information intact, however, it also misses neighboring information when unpooling from low-resolution feature maps. Finally, a Softmax classifier is used to produce the final segmentation maps with the same resolution of the original MG image. In this work, we used a Segnet that is pre-initialized with layers and weights from a pre-trained VGG16 model with an encoder D of 5.

#### Semantic segmentation using Dilated-Net

Recently, Dilated-Net [9], also known as atrous convolutions, have been used in different image segmentation tasks [10–14]. Dilated convolutions [9] allow us to explicitly control the resolution at which feature responses are computed and incorporate larger context without increasing the number of parameters or the amount of computation. We adopt the dilated CNN in [9] with some modification to the network. The implemented dilated CNN architecture consists of ten cascaded 3×3 convolutional layers with dilation factors 1, 1, 2, 4, 8, 16, 32, 1, 1 and 1 (Fig. 4). Figure 4 illustrates a 3×3 convolution kernels with different dilation factor as 1,2, and 3. The last three layers are FCLs of 1×1 convolutions followed by dropout of 0.5 [15]. The first nine convolutional layers are followed by BN layer [16] and a ReLU activation function [17]. To classify the pixels, the last convolutional layer has two 1×1 convolutions, followed by a Softmax classifier.

#### Localization using Faster R-CNN

We adapt the Faster R-CNN method proposed in [18] to compare its performance in terms of accuracy of detection and inference time with that of the proposed Vanilla U-Net model. Faster R-CNN is based on a VGG16 model [7] with additional components for detecting, localizing and classifying lesions in MG image. Faster R-CNN outputs a BB for each detected lesion, and a score, which reflects the confidence in the class of the lesion. The Faster R-CNN method in [18] is trained with our pre-processed, and augmented data-set. Further details about the implemented Faster R-CNN method can be found in the original article in [18]. One limitation stated in the study of [18] is that the training data-set comes from small sized publicly available pixel-level annotated data-set. However, in our study we are using our combined large-sized data-set to reproduce their work.

#### Semantic segmentation using region growing (RG)

We also implemented the region growing (RG) model proposed in [19] and apply it to our MG images. RG is a traditional image segmentation CAD model that starts with selecting an initial seed point and then groups pixels or sub-regions into larger regions according to a similarity criterion. As RG results are sensitive to the initial seeds, the automated accurate seed selection is very critical for image segmentation. Further details about the implemented RG method can be found in the original article [19].

#### Comparison between state-of-the-art DL methods

Table 1 lists the information about the architecture, databases, the number of images, the evaluation methods (i.e. Accuracy (ACC.), area under curve (AUC), Dice index (DI)), TPR@FPR, the testing time per image as provided in the literature.

**Author details**

#### List of tables

Table 1 Shows a comparison between the proposed segmentation method and the current state-of-the-art DL methods for segmentation or localization of lesions in MG images.

#### List of figures

Fig. 1 The databases used in our experiments.
Fig. 2 MG images and their corresponding GTMs.
Fig. 3 In Segnet, the indices at each max-pooling layer in the encoder contracting path at each level are stored and later used to up-sample the corresponding feature map in the decoder by unpooling it using those stored indices.
Fig. 4 Architecture of the dilated-Net, containing ten convolutional layers with dilation factors, indicated in red, increasing from 1 in the first layer to 32 in the seventh layer. The last 1×1 convolutional layer is followed by a Softmax classifier.

**Table 1**

| Paper | Approach | Tested on | #Images | Size | TL | Aug. | AUC | ACC. | DI | TPR@FPR | Inference time/image |
|---|---|---|---|---|---|---|---|---|---|---|---|
| **Proposed method** | Vanilla U-Net | INbreast, DDSM, UCHCDM | 17,140 | 512×512 | Yes | Yes | - | 0.926 | 0.951 | - | 0.12 sec |
| **Rilbi et al. [18]** | Faster R-CNN | INbreast | 3,927 | 1400×1700 | Yes | Yes | 95.0 | N/A | N/A | 0.90@0.3 | N/A |
| **Choukroun et al. [20]** | Batch-based CNN | INbreast | 410 | 224×224 | Yes | Yes | 72.2 | N/A | N/A | N/A | N/A |
| | | INbreast | 2,500 | 224×224 | Yes | Yes | 83.1 | N/A | N/A | N/A | N/A |
| **Akselrod et al. [21]** | Faster R-CNN | In-house | 850 | 800×800 | Yes | Yes | 72.0 | 77.0 | N/A | 0.93@0.56 | 0.2 sec |
| **Akselrod et al. [22]** | Faster R-CNN | INbreast<br>INbreast | 100<br>310 | 224×224<br>224×224 | Yes<br>Yes | Yes<br>Yes | N/A<br>N/A | N/A<br>N/A | N/A<br>N/A | 0.93@0.56<br>0.8@1.5 | 5 sec<br>5 sec |
| **Dhungel et al. [23]** | R-CNN+ Random forest (RF) | INbreast | 410 | 40×40 | Yes | Yes | N/A | 95.0 | N/A | 0.91@1.3 | 41 sec |
| **Dhungel et al. [3]** | R-CNN+ RF | INbreast | 410 | 40×40 | Yes | Yes | 91.0 | 95.0 | 85 | 0.9@1.0 | 41 sec |
| **Platania et al. [24]** | YOLO | DDSM | 10,480 | 128×128 | Yes | Yes | N/A | 90.0 | N/A | N/A | 1.25 sec |
| **Teuwen et al. [25]** | Faster R-CNN<br>Mask R-CNN | In-house<br>In-house | 23,405<br>23,405 | 227×227<br>227×227 | Yes<br>Yes | Yes<br>Yes | N/A<br>N/A | N/A<br>N/A | N/A<br>N/A | 0.87@0.56<br>0.97@3.5 | N/A<br>N/A |
| **Al-masni et al. [26]** | YOLO | DDSM | 600 | 448×448 | No | No | 87.74 | 96.33 | N/A | 0.96@2.6 | N/A |
| **Al-masni et al. [27]** | YOLO | DDSM | 600 | 448×448 | No | Yes | 96.45 | 99.70 | N/A | N/A | 3 sec |
| **Sun et al. [28]** | Batch-based CNN | In-house | 840 | 52×52 | No | Yes | 69.82 | N/A | N/A | N/A | N/A |
| **Reiazi et al. [29]** | Faster R-CNN | MIAS | 120 | 75×75 | No | No | N/A | N/A | N/A | N/A | N/A |

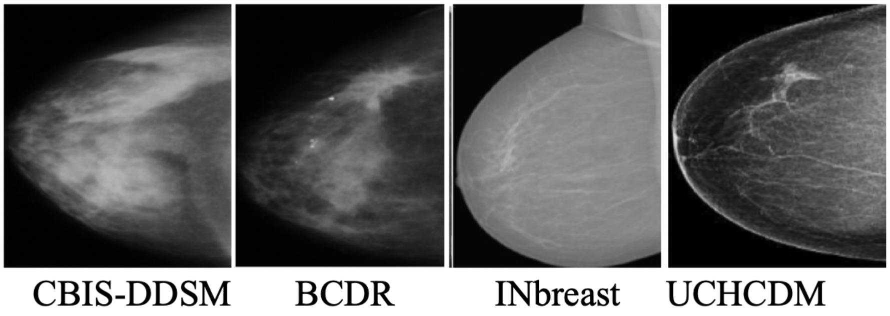


**Fig. 1**

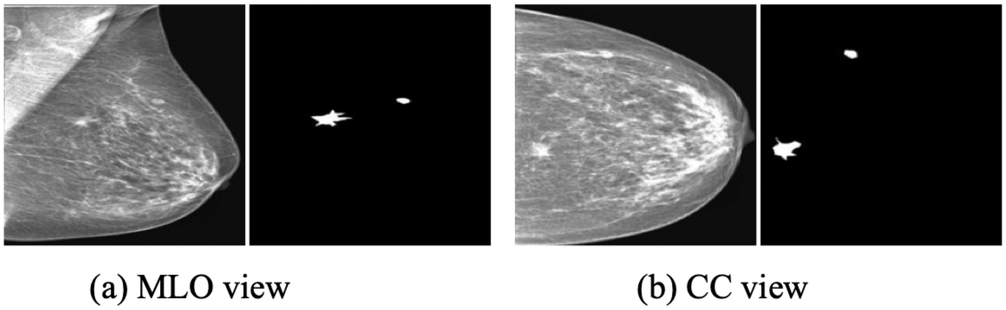
(a) MLO view
(b) CC view

Fig. 2

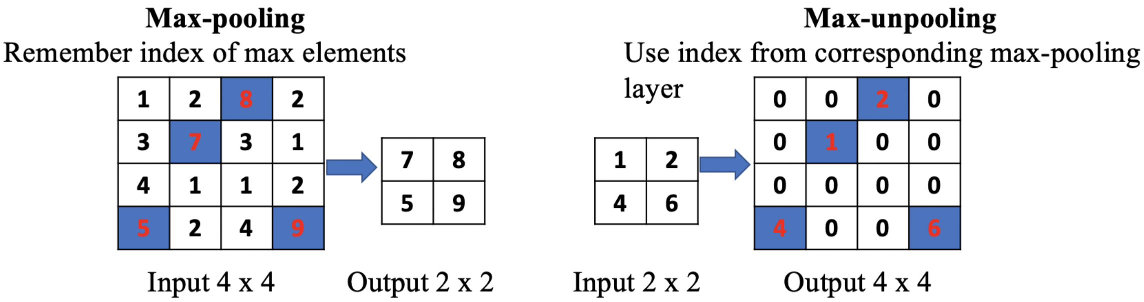
Max-pooling
Remember index of max elements
1 2 8 2
3 7 3 1
4 1 1 2
5 2 4 9
7 8
5 9
Input 4 x 4
Output 2 x 2
Max-unpooling
Use index from corresponding max-pooling layer
1 2
4 6
0 0 2 0
0 1 0 0
0 0 0 0
4 0 0 6
Input 2 x 2
Output 4 x 4

Fig. 3

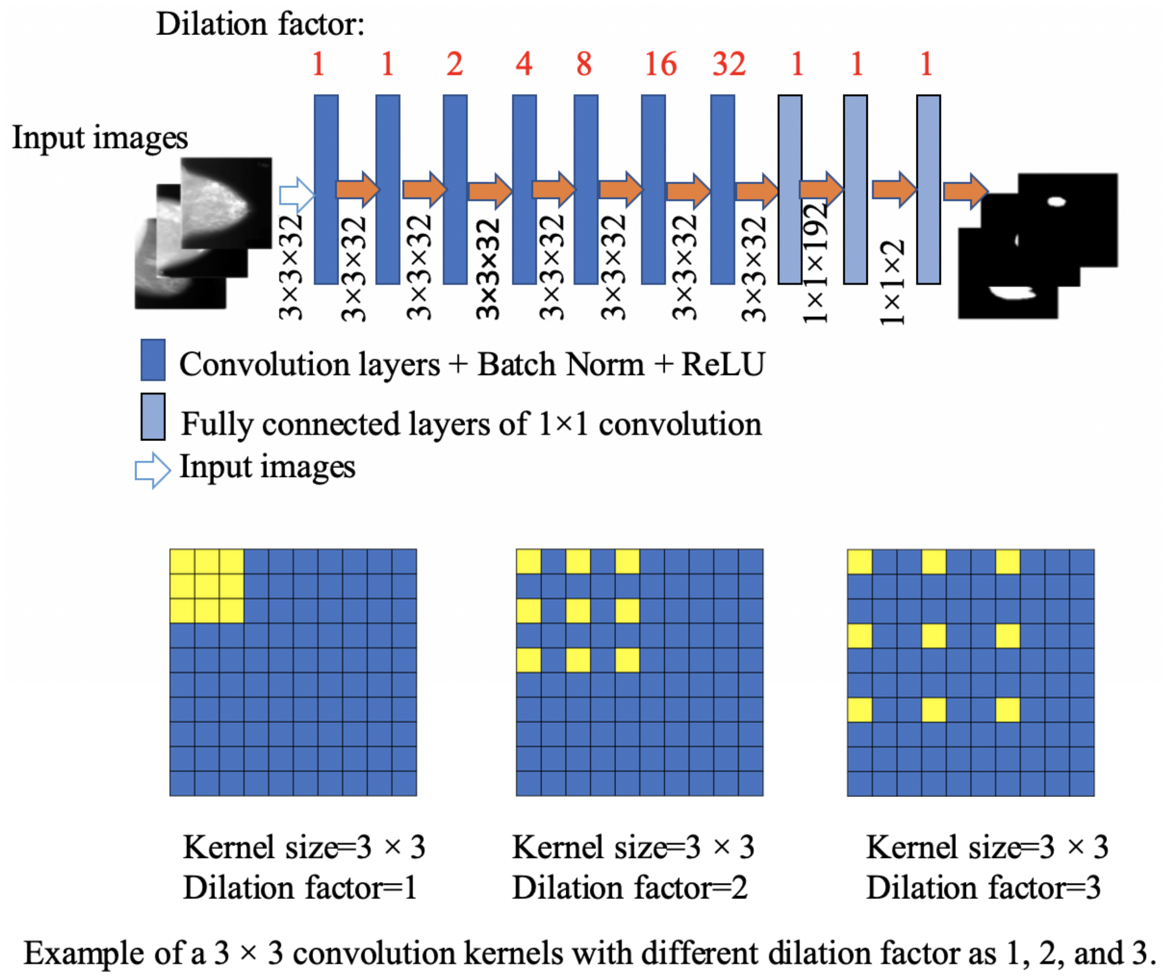
Dilation factor:
1 1 2 4 8 16 32 1 1 1
Input images
3×3×32
3×3×32
3×3×32
3×3×32
3×3×32
3×3×32
3×3×32
3×3×32
1×1×192
1×1×2
Convolution layers + Batch Norm + ReLU
Fully connected layers of 1×1 convolution
Input images
Kernel size=3 × 3
Dilation factor=1
Kernel size=3 × 3
Dilation factor=2
Kernel size=3 × 3
Dilation factor=3
Example of a 3 × 3 convolution kernels with different dilation factor as 1, 2, and 3.

**Fig. 4**
